# Transcribed germline-limited coding sequences in *Oxytricha trifallax*

**DOI:** 10.1101/2020.10.07.330092

**Authors:** Richard V. Miller, Rafik Neme, Derek M. Clay, Jananan S. Pathmanathan, Michael W. Lu, V. Talya Yerlici, Jaspreet S. Khurana, Laura F. Landweber

**Affiliations:** Department of Biochemistry and Molecular Biophysics, Columbia University, New York, NY 10032, USA; Department of Molecular Biology, Princeton University, Princeton, NJ 08544, USA; Department of Chemistry and Biology, Universidad del Norte, Barranquilla, Colombia; School of Environmental and Biological Sciences, Rutgers University, New Brunswick, NJ 08901, USA; Department of Laboratory Medicine and Pathobiology, Faculty of Medicine, University of Toronto, Toronto, ON M5G 1M1, Canada; Strand Therapeutics, Cambridge, MA 02139, USA; Department of Biological Sciences, Columbia University, New York, NY 10027, USA

**Keywords:** germline, genome rearrangement, DNA elimination, noncoding RNA, ciliate, micronucleus

## Abstract

The germline-soma divide is a fundamental distinction in developmental biology, and different genes are expressed in germline and somatic cells throughout metazoan life cycles. Ciliates, a group of microbial eukaryotes, exhibit germline-somatic nuclear dimorphism within a single cell with two different genomes. The ciliate *Oxytricha trifallax* undergoes massive RNA-guided DNA elimination and genome rearrangement to produce a new somatic macronucleus (MAC) from a copy of the germline micronucleus (MIC). This process eliminates noncoding DNA sequences that interrupt genes and also deletes hundreds of germline-limited open reading frames (ORFs) that are transcribed during genome rearrangement. Here, we update the set of transcribed germline-limited ORFs (TGLOs) in *O. trifallax*. We show that TGLOs tend to be expressed during nuclear development and then are absent from the somatic MAC. We also demonstrate that exposure to synthetic RNA can reprogram TGLO retention in the somatic MAC and that TGLO retention leads to transcription outside the normal developmental program. These data suggest that TGLOs represent a group of developmentally regulated protein coding sequences whose gene expression is terminated by DNA elimination.

## Introduction

Ciliates are a lineage of microbial eukaryotes characterized by functional nuclear differentiation. Each ciliate cell has one or more somatic macronuclei (MAC) and one or more germline micronuclei (MIC). The somatic MAC contains the somatic genome, consisting of over 17,000 gene-sized nanochromosomes that are transcribed throughout the organism’s life cycles (Swart et al. 2013; Lindblad et al. 2019). The germline genome is a fragmented and scrambled version of the somatic genome that undergoes a complex process of DNA deletion and rearrangement during sexual reproduction (Chen et al. 2014).

Previous studies have shown that *Oxytricha*’s sexual rearrangement cycle is guided by several noncoding RNA pathways. In the early stages of the sexual life cycle, bidirectional transcription across the length of nanochromosomes produce thousands of long template RNAs from the parental MAC (Lindblad et al. 2017). These transcripts guide the rearrangement of macronuclear destined sequences (MDSs) during development, and previous experiments showed that injection of synthetic template RNAs could program aberrant rearrangements (Nowacki et al. 2008; Bracht et al. 2017; Nowacki et al. 2011). Millions of 27-nucleotide long PIWI-associated small RNAs (piRNAs) are abundant during early *Oxytricha* rearrangement and interact with the *Oxytricha* PIWI ortholog Otiwi-1. These piRNAs also derive from the parental MAC. Their role is to protect the sequences they target against DNA deletion during development of the zygotic MAC. Injection of synthetic piRNA sequences that target internal eliminated sequences (IESs) that interrupt MDSs in the MIC can prevent their deletion during rearrangement and program their retention in the MAC (Fang et al. 2012). Programmed IES retention is now used as a genetic tool to create somatic knockout strains in *Oxytricha* (Khurana et al. 2018; Beh et al. 2019).

Besides IESs and transposons that are eliminated during development, *Oxytricha* has other classes of germline-specific MIC DNA sequences (Chen et al. 2016). Analysis of the germline MIC genome together with transcriptome-guided gene prediction previously uncovered 810 germline-limited protein coding genes encoded in the MIC genome (Chen et al. 2014). These germline-limited genes are specifically transcribed during rearrangement, and 26% of them had demonstrated translation of peptides present in a survey of one developmental time point.

Other lineages also have germline-limited protein coding sequences, including the ciliate *Tetrahymena thermophila* (Hamilton et al. 2016; Lin et al. 2016; Feng et al. 2017), the parasitic roundworm *Ascaris suum* (Wang et al. 2012; Wang et al. 2017), and the sea lamprey *Petromyzon marinus* (Bryant et al. 2016; Smith et al. 2009; Smith et al. 2012; Timoshevskiy et al. 2016; Timoshevskiy et al. 2017). Protein coding sequences are discarded in all these cases, and genes eliminated from somatic lineage cells are typically predicted to have functions in the germline and embryogenesis (Smith et al. 2012; Bryant et al. 2016). The songbird *Taeniopygia guttata* has a germline-limited chromosome that is deleted from somatic lineage cells (Pigozzi and Solari 1998; Pigozzi and Solari 2005; Itoh et al. 2009; Biederman et al. 2018; Kinsella et al. 2019; Torgasheva et al. 2019).

Here, we update and expand the set of transcribed germline-limited ORFs (TGLOs) in *Oxytricha* and provide functional experiments that reprogram the somatic retention of a small number of TGLOs to test the hypothesis that developmental deletion is the main mechanism to repress their gene expression during asexual growth. Like the previous set of germline-limited genes, we show that TGLOs contain several predicted functions and conserved domains that could be involved in Oxytricha development. This work also identified a locus, g111288, that is retained in the somatic MAC of a subset of progeny cells, revealing an example of a strain-specific macronuclear chromosome.

## Materials and methods

### Illumina library preparation and sequencing

Genomic DNA was collected from mated *O. trifallax* cells at various developmental time-points using the Nucleospin genomic DNA spin column column (Machery-Nagle). Illumina DNA sequencing libraries were prepared using the NEBNext Ultra II library preparation kit (New England Biolabs). 2 × 250 bp paired end sequencing reads were obtained using an Illumina HiSeq 2500, and remaining adapter sequences were trimmed using Trim Galore! software in the Galaxy cloud computing environment.

Total RNA was extracted from mated *O. trifallax* cells at various developmental time-points using Trizol reagent (Thermo Fisher, Waltham, MA, USA). Contaminating DNA was removed using a Turbo DNase kit (Thermo Fisher, Waltham, MA, USA). Poly-adenylated transcripts were enriched using the NEBNext Poly(A) mRNA Magnetic Isolation Module (New England Biolabs, Ipswich, MA, USA). RNA sequencing libraries were prepared using the ScriptSeq version 2 kit (Illumina, San Diego, CA, USA). 2 × 75 bp paired end sequencing reads were obtained using an Illumina HiSeq 2500, and remaining adapter sequences were trimmed using Trim Galore! software in the Galaxy cloud computing environment.

### TGLO computational prediction

We predicted TGLOs using a previously published pipeline for germline-limited gene prediction with some modifications (Chen et al. 2014). We predicted coding sequences with AUGUSTUS (version 3.3.0) (Stanke et al. 2006) using a gene prediction model trained on *O. trifallax* somatic MAC genes and transcripts as hints. We generated hint files for the gene prediction software by mapping RNA-seq data from cells collected at various time points to the germline MIC genome using HISAT2 (version 2.0.5). We ran AUGUSTUS with the options -- UTR=on and --alternatives-from-evidence=true. We filtered AUGUSTUS gene predictions to keep only models supported by hints including at least four supporting RNA-seq reads and greater than 80% of the coding sequence covered by RNA-seq reads to obtain the high transcription dataset. We kept only models supported by hints including at least two supporting RNA-seq reads and required greater than 20% of the coding sequence be covered by RNA-seq reads to obtain the low transcription dataset. We also removed candidate sequences with more than a minimal number of whole cell genomic DNA reads mapped from asexually growing cultures of either parental genotype or a pool of F1 cells to ensure that MAC encoded candidates were removed while accounting for the fact that some MIC encoded sequences will be present in whole cell sequencing reads.

### DNA sequencing analysis

Genomic DNA sequencing reads were aligned to the *O. trifallax* MIC genome assembly using BWA-MEM (version 0.7.17) with the -M option to mark short split alignments as supplementary alignments. Alignment files were processed using the Samtools software package (version 0.1.20) (Li et al. 2009). FeatureCounts software (version 2.0.0) (Liao et al. 2014) was used to assess the raw number of reads mapping to *O. trifallax* genome features (Burns et al. 2016). Relative DNA copy number changes for each genome feature were normalized using the R/Bioconductor package DESeq2 (version 1.26.0) (Love et al. 2014). Heat maps showing normalized DNA copy number during the developmental life cycle were generated using the log2 normalized copy number values and the pheatmap R package (version 1.0.12).

### Transcriptome sequencing analysis

Poly(A)-selected RNA sequencing reads were aligned to the *O. trifallax* MAC genome assembly and MIC genome assembly using HISAT2 (version 2.0.4) and Bowtie2 in the local alignment mode, respectively. Relative DNA copy number changes were normalized using the R/Bioconductor package DESeq2. Alignment files were processed using the Samtools software package (version 0.1.20) (Li et al. 2009). FeatureCounts software (version 2.0.0) (Liao et al. 2014) was used to assess the raw number of reads mapping to *O. trifallax* genome features (Burns et al. 2016). Relative RNA expression changes for each genome feature were normalized using the R/Bioconductor package DESeq2 (version 1.26.0) (Love et al. 2014). Heat maps showing normalized RNA expression during the developmental life cycle were generated using the log2 normalized copy number values and the pheatmap R package (version 1.0.12). Two timepoints of triplicate RNA-seq reads (12 hr and 36 hr) from the late time-course were previously uploaded to the European Nucleotide Archive under the project number PRJEB32087.

### Small RNA sequencing analysis

Previously sequenced Otiwi-1-dependent piRNAs (Fang et al. 2012) were aligned to the

*O. trifallax* MIC genome assembly using Bowtie2 (version 2.3.4.1) in the local alignment mode. Alignment files were processed using the Samtools software package (version 0.1.20) (Li et al. 2009), and alignments were viewed in the context of the MIC genome using the Integrative Genomics Viewer (version 2.7.2) (Robinson et al. 2011).

### Mass spectrometry analysis

Raw data were analyzed using MaxQuant (version 1.6.3.4) to search against a combined database containing previously published macronuclear-encoded and MIC-limited genes in addition to either highly-transcribed or lowly-transcribed TGLOs (Chen et al. 2014). Searches were performed using Trypsin/P as the enzyme with a maximum of two missed cleavages, methionine oxidation and protein N-terminal acetylation as variable modifications, cysteine carbamidomethylation as a fixed modification, precursor mass tolerances of 20 ppm for the first search and 4.5 ppm for the main search, and a maximum FDR of 1% for both peptides and proteins.

### Cell culture

*Oxytricha trifallax* cells were cultured in Petri dishes or large Pyrex dishes containing Pringsheim medium (0.11 mM Na_2_HPO_4_, 0.08mM MgSO_4_, 0.85 mM Ca(NO_3_)_2_, 0.35 mM KCl, pH 7.0) and fed *Chlamydomonas reinhardtii* and *Klebsiella pneumoniae* according to previously published methods (Khurana et al. 2014). Matings were performed by starving the compatible parental mating types 310 and 510, mixing the mating types, and diluting to a concentration of about 5,000 cells per milliliter in Pringsheim medium and plating the cells in 10 cm plastic Petri dishes. Matings were assessed several hours after mixing mating types by calculating the percentage of paired cells per total cells.

### Reverse transcription PCR (RT-PCR)

Cell cultures or mating time-courses were concentrated by centrifugation and total RNA was extracted using Trizol. Turbo DNase (Thermo Fisher, Waltham, MA, USA) was used to digest DNA before extracting RNA again. Eluted DNA-free total RNA was reverse transcribed using oligo (dT) and AMV reverse transcriptase (New England Biolabs, Ipswich, MA, USA). PCR was performed using cDNA template and Phusion High Fidelity DNA polymerase (New England Biolabs, Ipswich, MA, USA).

### Nanochromosome assembly

Pooled F1 cells were sequenced using Illumina sequencing. Short reads were mapped to the germline MIC genome. Reads mapping to g111288 were isolated. Next, we searched for the 5’ and 3’ end of an arbitrary read mapping to g111288 in the other reads. We iterated the process of searching for the 5’ or 3’ end of each read in the remaining reads until we found a read terminating with a telomere repeat (C_4_A_4_). We manually assembled the sequences of the reads into an g111288 nanochromosome.

### In vitro transcription

To prepare long single-stranded RNA (ssRNA) transcripts for microinjection, PCR primers were first designed to use Phusion High-Fidelity DNA polymerase (New England Biolabs, Ipswich, MA, USA) to amplify the coding sequence of the desired TGLO and add a T7 promoter to the gene. The T7-flanked product was cloned using the TOPO TA cloning kit (Thermo Fisher, Waltham, MA, USA) and Sanger sequenced (Genewiz, South Plainfield, NJ, USA) to verify the insert. In vitro transcription was done using the HiScribe T7 High Yield RNA Synthesis Kit according to the manufacturer’s instructions (New England Biolabs, Ipswich, MA, USA).

### RNA injection

In vitro transcribed RNA was extracted using Trizol and resuspended to a concentration of 3 micrograms per microliter. ssRNA was microinjected into mating cells at 12 hours post-mixing according to previously published protocols (Fang et al. 2012). Post-injected cells were allowed to recover in Volvic water for two days before picking single cells and plating them in Volvic to establish clonal lines.

### 5’ rapid amplification of cDNA ends (5’ RACE)

We used a published 5’ RACE protocol (Scotto-Lavino et al. 2006) with minor changes. Briefly, total RNA was extracted in Trizol (Thermo Fisher, Waltham, MA, USA) and treated with Turbo DNase (Ambion). One microgram of DNase-treated total RNA was reverse transcribed using AMV reverse transcriptase (New England Biolabs, Ipswich, MA, USA) and a gene-specific primer for either the germline-limited gene or actin II control. cDNA was poly(A) tailed using terminal transferase (New England Biolabs, Ipswich, MA, USA). The A-tailed cDNA was amplified using two rounds of PCR amplification using Phusion High-Fidelity DNA Polymerase (New England Biolabs, Ipswich, MA, USA). The first round of amplification was done over 15 cycles, the first round product was diluted 1:1000, the diluted first round product was amplified over 35 cycles, and the products from the second round of amplification were resolved on an agarose gel and stained with ethidium bromide (Bio-Rad, Hercules, CA).

### RT-qPCR

As we did previously, we reverse transcribed total RNA from two different times during the organism’s life cycle using random hexamer primers. This cDNA was used as template in a series of RT-qPCR experiments to detect the expression of either germline-limited ORF candidate or actin. We used Power Sybr Green qPCR Master Mix (Thermo Fisher, Waltham, MA, USA) and custom qPCR primers (Integrated DNA Technologies, Coralville, IA, USA) and performed the reaction using a CFX384 Touch Real-Time PCR Detection System (Bio-Rad, Hercules, CA, USA). We analyzed the Cq values using a standard curve method and compared the number of transcripts in each sample to the number of small subunit mitochondrial rRNA.

### Southern hybridization

1 μg of genomic DNA was resolved on a 1% agarose gel, and ethidium bromide was used for visualization. MAC DNA was purified according to previously published methods (Swart et al. 2013). Dilute PCR products were used as a control to approximate the expected copy number in the genomic DNA lanes. The 1 Kb Plus DNA ladder (Thermo Fisher, Waltham, MA, USA) was used as a size standard. After gel electrophoresis, DNA was blotted onto a nylon membrane, detected using a digoxigenin-labeled DNA probe, and detected using chemiluminescence according to a previously published protocol (Yerlici et al. 2019)

### Primers

The following primers were synthesized by Integrated DNA Technologies (Coralville, IA, USA) for use in this study.

g104149 retention fwd: 5’-CGATGATGATGCAGAGCAGTGGAGGCTTAG-3’

g104149 retention rev: 5’-CATATCGTGTTCATTCATGTAAGATAACTACTGCTTG-3’

g67186 retention fwd: 5’-CAATTCACATAATCCTCTATTTCTGCAACTTTTTCTAGAC-3’

g67186 retention rev: 5’-GAATTATTTGTAAATACTTGACTGACTCATTGTTGATAAAATGATTTAC-3’

QT RACE: 5’-CCAGTGAGCAGAGTGACGAGGACTCGAGCTCAAGC-3’ (Scotto-Lavino 2006)

QO RACE: 5’-CCAGTGAGCAGAGTGACG-3’ (Scotto-Lavino 2006)

QI RACE: 5’-GAGGACTCGAGCTCAAGC-3’ (Scotto-Lavino 2006)

Actin II RT: 5’-GTGGTGAAGTTATATCCTCTCTTGGCCAATAATG-3’

Actin II GSP 1: 5’-TGGCATGAGGAATTGCGTAACCTTCATAGA-3’

Actin II GSP 2: 5’-TCCATCTCCAGAGTCAAGCACAACACC-3’

g104149 RT: 5’-TTGGGTAAATTCTGGCCAACTCCCTTG-3’

g104149 GSP 1: 5’-CCAAGCTTCTCTGCACCTCATCCGTGAACA-3’

g104149 GSP 2: 5’-GTCTGCCCATCCACGATTTCACTGACC-3’

g67186 RT: 5’-AGCCTTGGTCCCTTCTGAGGCAG-3’

g67186 GSP 1: 5’-CCTGGCAAGAGCAACTTGACAGCAC-3’

g67186 GSP 2: 5’-GAGAGGCCAGAGGCTTCATTGCATACC-3’

g104149 gene qPCR fwd: 5’-CCAAGCTTCTCTGCACCTCATCCGTGAACA-3’

g104149 gene qPCR rev: 5’-AAGGTCAGTGAAATCGTGGATGGGCAGACT-3’

g67186 gene qPCR fwd: 5’-TGCAATGAAGCCTCTGGCCTCTCA-3’

g67186 gene qPCR rev: 5’-CCTGGCAAGAGCAACTTGACAGCAC-3’

g67186 upstream qPCR fwd: 5’-CAATTCAATAGCACCGAATAGAAAGCTTATTTTATACAAGGATTAG-3’

g67186 upstream qPCR fwd: 5’-CTAGATTTAATTAAAACTTGAAATGTCTACAGCCCATTAATAATTCG-3’

Actin II qPCR fwd: 5’-GGTGTTGTGCTTGACTCTGGAGATGGA-3’

Actin II qPCR rev: 5’-TGGCATGAGGAATTGCGTAACCTTCATAGA-3’

Mitochondrial 23S rDNA qPCR fwd: 5’-GATAGGGACCGAACTGTCTCACG-3’ (Nowacki et al. 2009)

Mitochondrial 23S rDNA qPCR rev: 5’-CATATCCTGGTTGTGAATAATCTTCCAAGGG-3’ (Nowacki et al. 2009)

Telomere primer 1: 5’-ACTATAGGGCACGCGTGGTCGACGGCCCGGGCTGGTCCCCAAAACCCCAAAACCCC AAAA-3’ (Nowacki et al. 2008)

Telomere primer 2: 5’-ACTATAGGGCACGCGTGGT-3’ (Nowacki et al. 2008)

g43073 TSP 1: 5’-GCCAGGTAGTTGCAAGCGCTCTCGAGAG-3’

g43073 TSP 2: 5’-GCTCAAAGTTTTAACTACTTGATTGAAGTGTAGATTTGGCAATC-3’

g104149 TSP 1: 5’-GTAAATTCTGGCCAACTCCCTTGAGTTCCAAGCTTC-3’

g104149 TSP 2: 5’-CAAAGTCTGCCCATCCACGATTTCACTGACCTTTG-3’

g93797 TSP 1: 5’-GCCCAATTCATATGCTGCTTCTTTGAGCCACTTG-3’

g93797 TSP 2: 5’-GATCTGGTTTTCACAGTTGAGGTAGTAGTAGTAG-3’

g111288 fwd PCR: 5’-CTCTACTCTCTTAGGTCTCCCTCTGCCATT-3’

g111288 rev PCR: 5’-AGCGGCCTGAAACTTTGTAAGGAGTAAGAT-3’

Actin II fwd PCR: 5’-GACTCAAATTATGTTTGAAGTCTTCAATGTACCTTGCC-3’

Actin II rev PCR: 5’-GTGGTGAAGTTATATCCTCTCTTGGCCAATAATG-3’

g111288 nanochromosome gene fwd qPCR: 5’-CAGGCCGCTTTAACTGCAACCATAGTTG-3’

g111288 nanochromosome gene rev qPCR: 5’-GGAAATTGAGCCAACTTTACAGTTAGAGCC-3’

g111288 nanochromosome MDS2 fwd qPCR: 5’-CTTTCCTACAAATCCCCTTAAATTTCCAGTCTTGTAC-3’

g111288 nanochromosome MDS2 rev qPCR: 5’-GTACCATGCTAGGATGTTATTGAAATCATAGAAGAC-3’

g111288 nanochromosome MDS4 fwd qPCR: 5’-CGTCAAATTCAGTAACTAGCTCAGGTACGTC-3’

g111288 nanochromosome MDS4 rev qPCR: 5’-CTACCCTCCCGAGGAAAATACCTGG-3’

g111288 nanochromosome MDS7 fwd qPCR: 5’-CTGAAATGGCTGTATCTATGGTTATTATAAAGAATTAGTG-3’

g111288 nanochromosome MDS7 rev qPCR: 5’-CAATCATCACTCTCCCTAACCGTACCTC-3’

g111288 nanochromosome IES6 fwd qPCR: 5’-GGGAAGTTATTTTATTATGAGTTTAGGTTGCATTCATTC-3’

g111288 nanochromosome IES6 rev qPCR: 5’-GAATGAAAATGAGTGAATTAAGAATTTTAATGAAGTATGATATAACATTC-3’

### Bioinformatic analyses

Short read DNA sequences were locally aligned to reference sequences using Bowtie 2 (Langmead and Salzberg 2012) or BWA-MEM. Short read RNA sequences were aligned to reference sequences using HISAT2 (Kim et al. 2019). Sanger sequencing DNA reads were aligned to reference sequences using the Geneious aligner in the Geneious software package (version 5.9) (Biomatters, Ltd., Auckland, New Zealand) with default parameters.

### Data availability

All cell stocks are available upon request. Illumina sequencing datasets were uploaded to the NCBI Short Read Archive under the BioProject PRJNA665991. The authors affirm that all data necessary for confirming the conclusions of the article are present within the manuscript and figures.

## Results

### Thousands of transcribed germline-limited open reading frames (TGLOs) are expressed during development

We examined potential germline-limited coding sequences in the *Oxytricha trifallax* MIC genome by searching for transcribed germline-limited open reading frames, which we refer to as TGLOs. We adapted a computational pipeline originally used to identify 810 germline-limited protein coding genes expressed during *Oxytricha trifallax* development (Figure 1A, left) (Chen et al. 2014). First, we used Augustus gene prediction (Stanke et al. 2006) and RNA sequencing hints from throughout the organism’s life cycle to predict 217,805 potential coding sequences in the germline genome. To exclude potential coding sequences that are present in the somatic MAC genome or are not transcribed at significant levels, we restrict TGLOs to computationally predicted ORFs with virtually no DNA sequencing coverage in the MAC genome of both parental strains. Another requirement is that they have RNA expression in at least one timepoint during the organism’s life cycle. To set read mapping thresholds appropriate for the variable sequencing depth of individual RNA and DNA libraries, we used a Monte Carlo approach in which the predicted 217,805 candidate loci were randomly shuffled 100 times throughout the germline-limited portion of the MIC genome, while recording the distribution of the number of DNA and RNA reads mapped to the random loci. The distributions of DNA or RNA reads mapped to randomly shuffled TGLO loci were treated as the background germline-limited coverage. We required that TGLOs have a number of DNA sequencing reads mapping to them from either parent or the F1 progeny that is no greater than the fifth percentile from the background germline-limited coverage simulation (i.e. no reads mapped per TGLO). On the other hand, highly expressed TGLOs should have RNA sequencing coverage equal to at least the 95th percentile from the random distribution (i.e. four reads mapped per TGLO). We also used a lower RNA sequencing threshold (i.e. a minimum of two reads mapped per TGLO) because at least one experimentally confirmed TGLO was not present in the high transcription TGLO dataset. CD-HIT (Fu et al. 2012) and RepeatMasker (Smit et al. 2013) were used to cluster similar sequences and to remove sequences associated with repetitive elements. The final mutually exclusive datasets contained 4342 and 6296 TGLOs with high and low transcription levels, respectively (Figure 1A, center). Like the previously reported germline-limited gene dataset, TGLOs tend to be intron-poor, with 8.8% and 6.4% of high and low transcription TGLOs, respectively, containing introns compared to 64.9% of MAC encoded genes. These datasets update our previous estimates and contain 279 (213 and 66, resp.) of the 810 germline-limited genes predicted in Chen et al. (2014) (Figure 1A, right) (Chen et al. 2014), with some of the reduction attributed to strain-specific differences described below.

**Figure 1:**
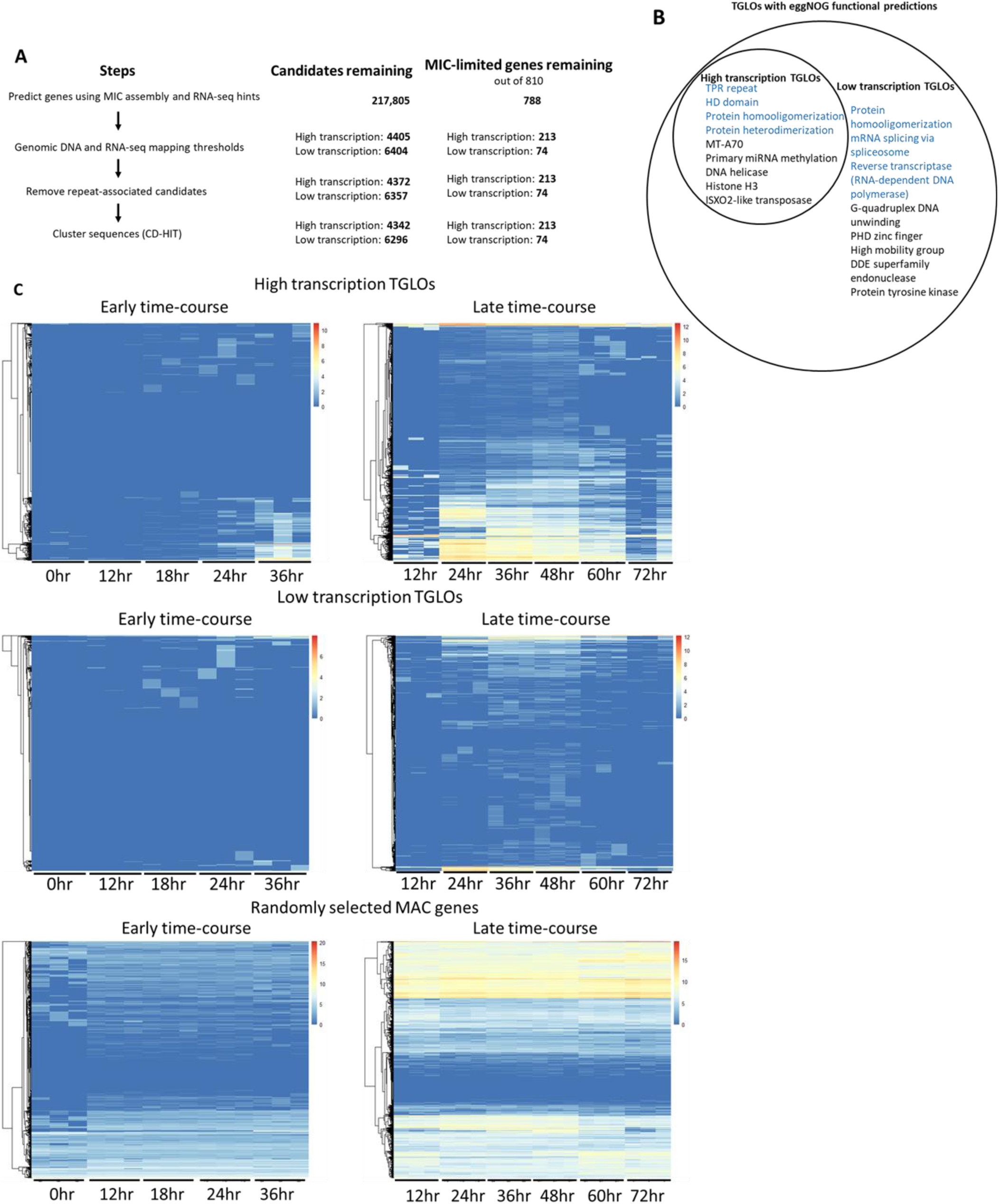
Germline-limited ORFs are expressed during *Oxytricha trifallax* genome rearrangement. **(A)** Left: Pipeline for predicting TGLOs in *Oxytricha trifallax* germline MIC genome. Center: Total number of computationally predicted candidates remaining after each pipeline step. Right: Total number of MIC-limited genes (Chen et al. 2014) remaining after each pipeline step. **(B)** EggNOG mapper-predicted functions and conserved domains in TGLOs. Blue text indicates that peptides from the associated TGLOs were present in a single nuclear proteome surveyed during rearrangement (Chen et al. 2014). **(C)** Log_2_-normalized RNA-seq read counts of High and Low transcription TGLOs and one thousand randomly selected somatic MAC-encoded genes across the *Oxytricha trifallax* developmental life cycle (hours labeled post mixing of compatible mating types). Color scale refers to the log2-normalized RNA expression.

The previous set of 810 germline-limited genes included functional predictions (Chen et al. 2014). We investigated conserved domains and putative gene functions using the functional annotation tool eggNOG mapper (version 2) (Huerta-Cepas et al. 2017). 111 high transcription TGLOs and 245 low transcription TGLOs mapped to conserved eggNOG orthology clusters (version 5.0) (Figure 1B) (Huerta-Cepas et al. 2019). 54 TGLOs with functional predictions were previously-predicted germline-limited genes (42 and 12 in high and low transcription TGLOs, respectively). Predicted functions and conserved domains included several potentially involved in DNA rearrangement and epigenetic regulation, including MT-A70, miRNA methylation, DNA helicase, PHD zinc finger, and high mobility group.

Protein expression of TGLOs could also suggest a function role for a subset of predicted coding sequences. One quarter (26%) of the original 810 germline-limited genes had peptides identified in a nuclear proteome extracted from mid-rearrangement cells at a single timepoint (Chen et al. 2014), and we queried the new TGLO datasets against this previously published peptide dataset. 144 high and 48 low transcription TGLOs (101 and 42 newly discovered, respectively) were present in this limited 40 hour proteomic survey. Several peptides from the developmental survey were also mapped to TGLOs with eggNOG functional predictions (Figure 1B, blue text).

The previously published set of germline-limited genes was limited to developmental gene expression, with most germline-limited genes transcribed beginning 40 hours after mixing of parental cells (Chen et al. 2014). We assessed the transcription profiles of TGLOs throughout the organism’s developmental life cycle using a deeply sequenced set of developmental RNA sequencing libraries. Two partially overlapping triplicate RNA sequencing time-courses across post-zygotic development showed that RNA expression from both high (Figure 1C) and low transcription TGLOs also clustered toward the later stages of rearrangement. Conversely, a random sample of one thousand somatic MAC-encoded genes had a diverse set of RNA expression profiles during the same time-course, suggesting that TGLOs are enriched in developmental expression.

### TGLO genes are eliminated after gene expression

By definition, TGLO DNA sequences are restricted to the germline MIC. Since the germline genome is diploid, TGLOs are present at a copy number equal to twice the number of micronuclei per cell. Since DNA copy number changes significantly throughout MAC development (Spear and Lauth 1976), we studied DNA copy number changes and elimination of TGLOs during development. A preliminary copy-number study indicated that most TGLOs are eliminated by the end of the developmental life cycle, but the DNA copy number profiles of TGLOs are heterogeneous, with some showing very little copy number variation throughout development, leaving it unclear whether the loci are eliminated from the developing somatic MAC by the end of the sexual life cycle (Figure 2A).

**Figure 2:**
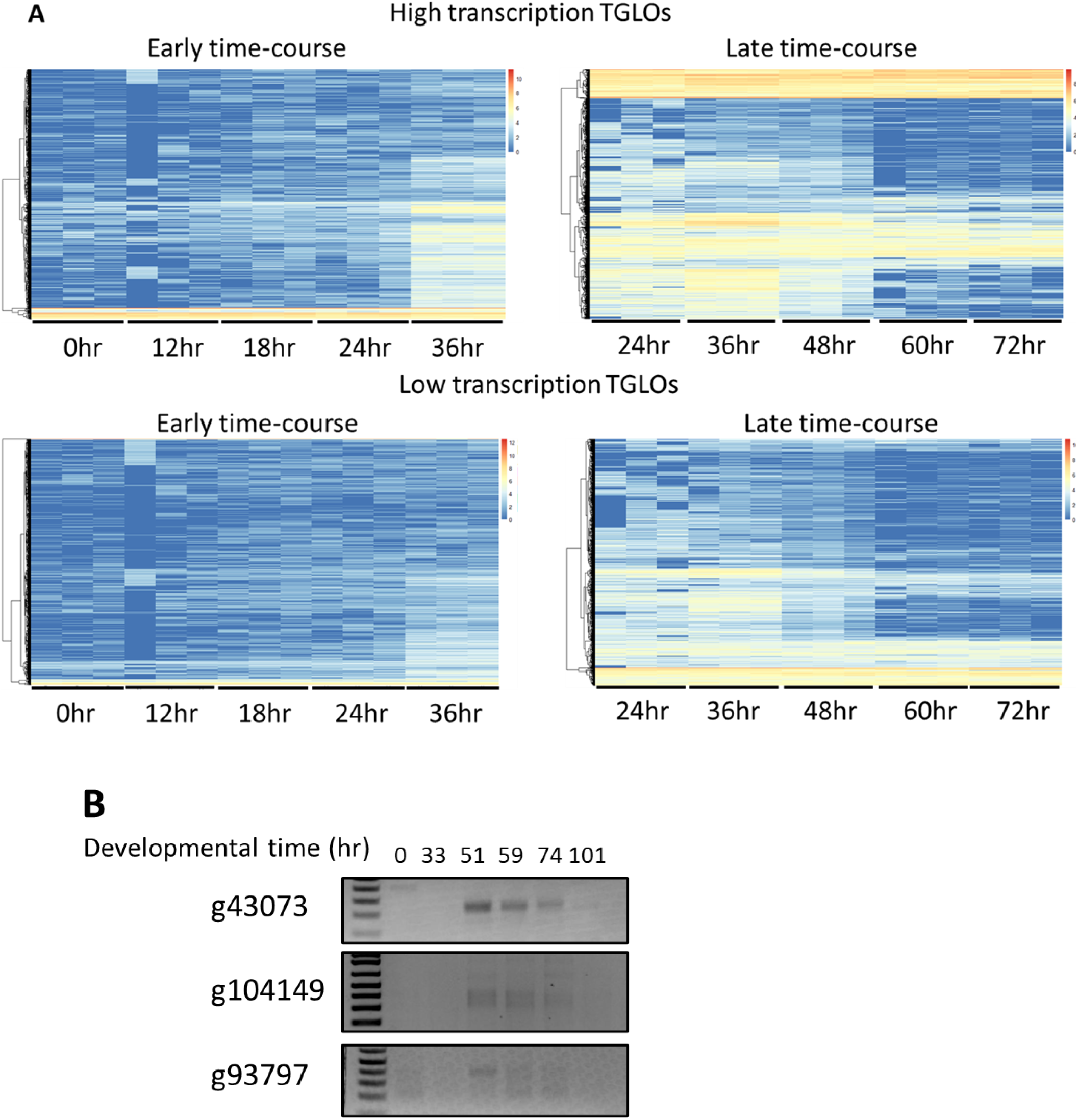
TGLOs are eliminated from the developing MAC. **(A)** Log_2_-normalized DNA copy number of High and Low transcription TGLOs across the *Oxytricha trifallax* developmental life cycle. Color scale refers to the Log_2_-normalized DNA copy number. **(B)** Telomere suppression PCR targeting the upstream telomere addition site of selected TGLOs in genomic DNA samples collected throughout the *Oxytricha trifallax* developmental life cycle.

Since we previously reported that telomeres are permissive to transcription in *O. trifallax*, unlike in other lineages (Beh et al. 2019), we amplified several TGLO loci via telomere suppression PCR (Chang et al. 2004) to determine whether telomeres are added upstream of these loci before DNA elimination. We found that three out of six sampled TGLOs—representing both high and low transcription TGLOs—had telomeres added near the ORF during mid to late development and before their elimination from the developing somatic MAC (Figure 2B), consistent with their transcriptional pattern.

### Strain-specific germline-limited ORFs

Our studies uncovered one case of a germline-encoded ORF that was also present at a low copy level in the somatic MAC of one parent. The protein coding locus, OXYTRIMIC_220 (“g111288”), was included in the previously reported set of 810 MIC-limited genes, but it does not encode any conserved functional domains nor was it detected in a developmental mass spectrometry survey (Chen et al. 2014). The initial Augustus gene prediction identified this ORF. However, it was later excluded from the pipeline after incorporating new DNA sequencing libraries from the parent strains and F1 progeny, which suggested that g111288 is present in the somatic MAC of at least one parental strain.

We used PCR to amplify g111288 from parental genomic DNA to test whether the locus is present in the somatic genome of either parent strain. We found that the coding sequence was abundant in strain JRB510, which was not the reference strain used for genome sequencing (Swart et al. 2013; Chen et al. 2014). In addition, we found that several cell lines derived from either single F1 progeny or genetically manipulated F1 lines also contained g111288 at detectable DNA copy levels (Figure 3A). In addition, the g111288 locus varied in DNA copy level in individual F1 lines derived from different parental crosses.

**Figure 3:**
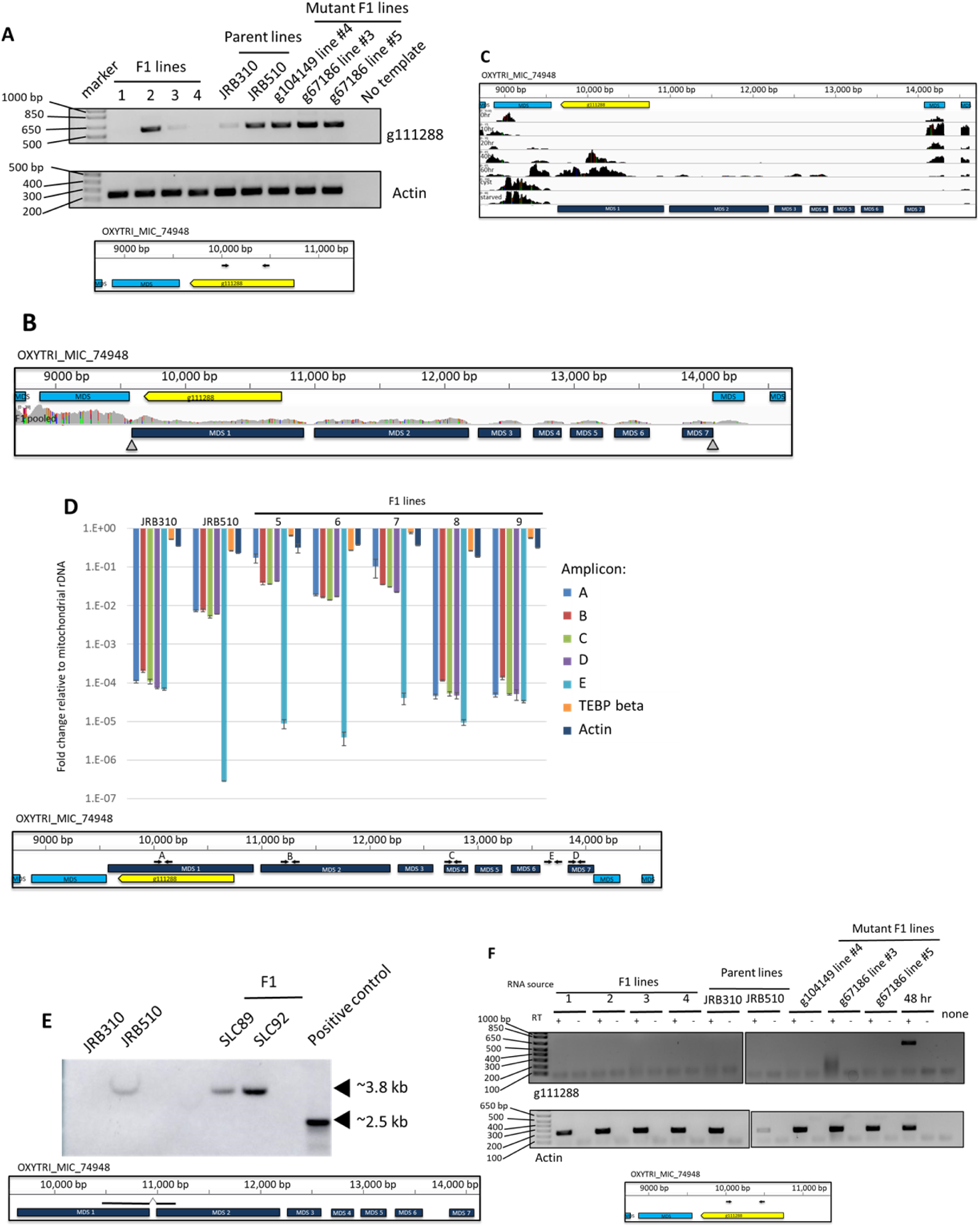
Parental cells can carry a strain-specific germline-limited ORF. **(A)** Top: PCR targeting g111288 or Actin II using genomic DNA from F1 lines, parent lines, and other mutant F1 lines used in this study. Bottom: Genome track showing the approximate location of g111288 PCR primers. Yellow: g111288, light blue: flanking MDSs. **(B)** The germline genome locus containing g111288 with mapped F1 reads from a pool of asexually growing F1 cells. Yellow: g111288, light blue: MDSs, dark blue: assembled g111288 MDSs from pooled F1 reads, gray triangles: observed telomere addition sites. **(C)** The germline genome locus (bottom) containing g111288 (yellow) and strain-specific MDSs (dark blue) with mapped RNA-seq coverage (black) from several time-points during asexual growth (starved or encysted cells) and hours post mixing of mating types during the sexual life cycle. **(D)** Top: Copy number relative to mitochondrial rDNA based on qPCR targeting several amplicons on the g111288 nanochromosome, an IES within the corresponding germline locus, and two unrelated somatic loci. Bottom: The germline genome locus containing g111288 with qPCR primer locations indicated. Yellow: g111288, light blue: MDSs, dark blue: assembled g111288 MDSs from pooled F1 reads, black arrows: qPCR primers. **(E)** Top: Southern blot of parental and F1 MAC DNA detected using an MDS-MDS junction spanning DNA probe. Bottom: MIC genome track showing the portions of MDSs 1 and 2 detected. **(F)** Top: RT-PCR targeting g111288 or Actin II using RNA from the same cell lines as in (A). Bottom: Genome track showing the approximate location of g111288 RT-PCR primers. Yellow: g111288, light blue: MDSs.

Since g111288 appeared to be present in the MAC genome of only parental strain, JRB510, and germline limited in the reference strain JRB310, we investigated the nature of the putative g111288 somatic MAC nanochromosome. Next generation sequencing reads from a pool of F1 progeny cells were mapped to the germline MIC genome. This allowed assembly of an entire g111288 nanochromosome with telomeres at both ends and indicated that it derives from seven MDSs with the g111288 open reading frame entirely contained within the first MDS (Figure 3B). RNA sequencing from developmental time-points confirmed that g111288 is transcribed from 40 to 60 hr after mixing of both parental strains (Figure 3C). In addition, alignment of RNA-seq reads to the other six MDSs on the g111288 nanochromosome suggested the possibility that the other six MDSs of the g111288 nanochromosome could have coding potential. To assess the nanochromosome’s relative copy number in different cell lines, we performed qPCR to target different amplicons across the g111288 nanochromosome using template genomic DNA from parental cells and F1 progeny lines. A two order of magnitude copy number increase was consistently observed in the JRB510 parent line relative to the reference JRB310 strain (Figure 3D). Moreover, three F1 lines displayed copy levels somewhat higher than the JRB510 parental strain, and the other two F1 lines appeared to have few to no copies of the nanochromosome, like strain JRB310. Southern hybridization with a probe targeting a MAC-specific MDS-MDS junction region confirmed the presence of the nanochromosome in MAC DNA from parental strain JRB510 as well as two F1 cell lines (SLC89 and SLC92; Seegmiller et al. 1996) (Figure 3E).

Since g111288 is present in the somatic genome of several F1 lines and at a low level in one parent, we assessed whether the coding sequence is transcribed during asexual (vegetative) growth. However, we did not detect any transcripts from this locus outside the middle and late stages of developmental, corresponding to approximately 48 hours after mixing of mating-compatible cells (Figure 3F). Swart et al. (2013) previously reported that many other MAC nanochromosomes have developmental-specific expression (Swart et al. 2013), suggesting that g111288 is a strain-specific nanochromosome, retained only in the MAC genome of JRB510 and passed on to its F1 progeny.

### Few ncRNAs map to TGLO loci

*Oxytricha*’s genome rearrangements and DNA deletion are regulated by noncoding RNAs (ncRNAs). For example, Otiwi-1-bound piRNAs map to retained MDSs but not germline-limited regions or IESs (Fang et al. 2012), and long template RNAs map to nanochromosomes in the MAC genome (Lindblad et al. 2017). Hence, we mapped template RNAs and Otiwi-1-associated piRNAs to the MIC genome and assessed their coverage in TGLO loci and the g111288 locus. We found that Otiwi-1 piRNAs map to MDSs more heavily than TGLOs (Figure 4A). Otiwi-1 piRNAs aligned to g111288, which is retained at a low somatic copy level in one parent (Figure 4B), but piRNAs are present at a reduced level compared to neighboring MDSs. Template RNA coverage was also significantly higher in MDSs compared to TGLOs (Figure 4C), although the strain-specific TGLO g111288 lacked any template RNAs despite being encoded by the JRB510 MAC (Figure 4D).

**Figure 4:**
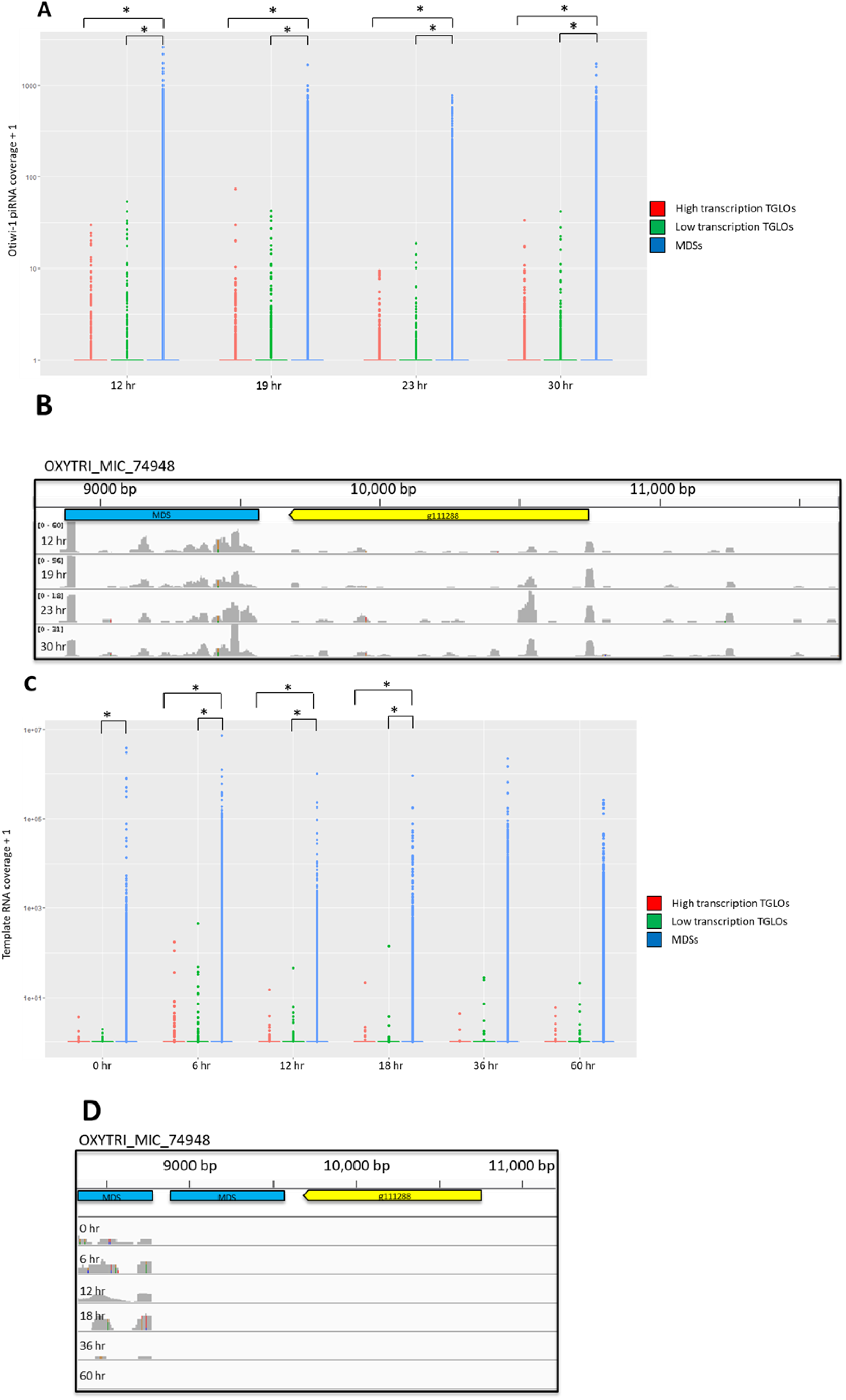
TGLO loci have few Otiwi-1 piRNAs and template RNAs. **(A)** Distribution of normalized mapping quality-filtered Otiwi-1 piRNA read counts (Fang et al. 2012) mapped to High and Low transcription TGLOs compared to MDSs. Read counts were normalized to reads per kilobase million (RPKM). Brackets and asterisks indicate statistically significant differences between distributions. Statistical significance was assessed using the non-parametric Kolmogorov–Smirnov (KS) test, and P<0.05 was considered statistically significant. **(B)** The germline genome locus containing the strain-specific TGLO g111288 (yellow), MDSs (blue), and mapped Otiwi-1-associated piRNA coverage (gray) from several time-points during rearrangement. **(C)** Distribution of normalized mapping quality-filtered template RNA read counts (Lindblad et al. 2017) mapped to High and Low transcription TGLOs compared to MDSs. Read counts were normalized to RPKM. Brackets and asterisks indicate statistically significant differences between distributions. Statistical significance was assessed using the non-parametric KS test, and P<0.05 was considered statistically significant. **(D)** The germline genome locus containing the strain-specific TGLO g111288 (yellow), MDSs (blue), and mapped template RNA coverage (gray) from several time-points during rearrangement.

### Synthetic RNA injection can protect TGLO loci from genomic deletion

g111288 presents an example of a potential coding sequence that is present in the somatic MAC of one strain while eliminated as a TGLO in another strain. We decided to test whether exposure to artificial RNAs during development could reprogram the germline-limited status of TGLOs, thereby retaining them on MAC nanochromosomes. Given our previous observations that exposure to non-coding RNAs can reprogram IES retention in the MAC (Fang et al. 2012; Khurana et al. 2018 RNA; Beh et al. 2019) we used RNA injection to test whether exposure to targeting RNA could reprogram the retention of two TGLO loci during development (Figure 5A). We targeted two TGLO loci that are encoded in the IESs of other MAC loci. One of the two candidates, g67186, was previously predicted to encode a histone 2B gene (Chen et al. 2014), while the other, g104149, did not contain any predicted conserved domains. The two candidates are also among the highest expressed TGLOs that mapped within IESs, facilitating our strategy (Figure 5A). Importantly, we also observed that our candidate TGLOs lacked Otiwi-1 piRNAs and template RNAs during the sexual life cycle (Figure 5B), suggesting that the cell does not endogenously encode their somatic retention during the sexual life cycle.

**Figure 5:**
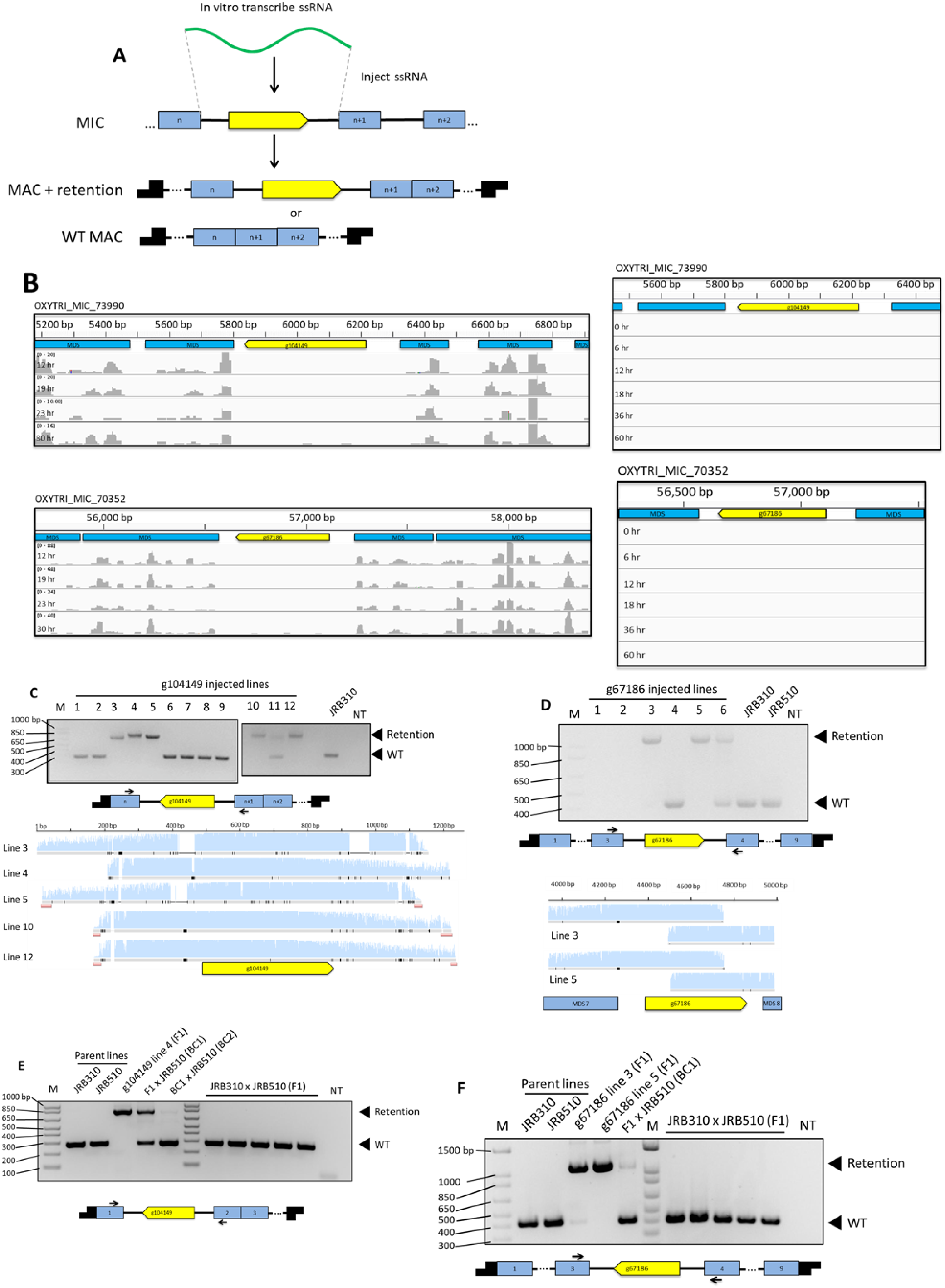
RNA injection programs heritable TGLO retention. **(A)** Synthetic RNA injection scheme to program the retention of a TGLO (yellow) in an IES between two MDSs (blue). Possible products can include telomere-capped (black) nanochromosomes with the entire IES plus TGLO flanked by the MDSs of the wild-type flanking locus. **(B)** The germline genome loci containing the programmed retention candidate TGLOs g104149 and g67186 (yellow), MDSs (blue), and mapped piRNA or template RNA coverage (gray) from several time-points during rearrangement. **(C)** Top: Cell culture PCR targeting the IES containing g104149 from cell lines derived from single RNA injected mating pairs. Middle: The expected retention product containing g104149 with PCR primer locations. Yellow: g104149, light blue: MDSs, black arrows: PCR primers. Bottom: Sanger sequencing chromatograms from PCR reactions in (B) aligned to the expected retention product containing g104149 (yellow). **(D)** Top: Cell culture PCR targeting the IES containing the predicted histone 2B TGLO g67186 from cell lines derived from single RNA injected mating pairs. Middle: The expected retention product containing g67186 with PCR primer locations. Yellow: g67186, light blue: MDSs, black arrows: PCR primers. Bottom: Sanger sequencing chromatograms from PCR reactions aligned to the expected retention product containing g67186 (yellow). **(E)** Top: PCR targeting the IES containing g104149 using genomic DNA from parental cells, F1 retention cells, F1 retention cells backcrossed to parental cells, and unmanipulated F1 lines. Bottom: The expected retention product containing g104149 with PCR primers. Yellow: g104149, light blue: MDSs, black arrows: PCR primers. **(F)** Top: PCR targeting the IES containing the predicted histone 2B TGLO g67186 using genomic DNA from parental cells, F1 retention cells, F1 retention cells backcrossed to parental cells, and unmanipulated F1 lines. Bottom: The expected retention product containing g67186 with PCR primers. Yellow: g67186, light blue: MDSs, black arrows: PCR primers.

PCR from cell cultures derived from single injected cells, followed by Sanger sequencing indicated that RNA injection did reprogram IES+TGLO retention in some progeny, with varying levels of retention based on differences in PCR band sizes. Some products contained small deletions in the retained sequence relative to the reference MIC locus, but no deletions affected the ORF (Figure 5C and 5D). No F1 lines from uninjected WT parental cells contained the TGLO sequences, suggesting that RNA injection specifically programs the somatic DNA retention (Figure 5E right and Figure 5F right).

RNA programmed IES retention was previously shown to be heritable after subsequent sexual cycles, so we also tested whether the IES+TGLO insertions were retained after backcrossing to a parental strain. PCR amplification from genomic DNA of backcrossed pools of cells indicated that the retained TGLO g104149 was partially heritable for at least two more generations (Figure 5E, left). The other retained TGLO, g67186, was partially heritable for one backcrossed generation (Figure 5F, left). A second band corresponding to the wild-type product was present in both backcrosses, consistent with the presence of WT nanochromosomes in the backcrosses to the wild-type parental strain.

### Retained TGLOs are transcribed outside usual developmental program

Our engineered strains that retain TGLO loci are unique in their ability to encode previously eliminated germline sequences in their macronucleus. Genome-wide transcription start site profiling in asexually growing *O. trifallax* cells showed that transcription initiation typically occurs in the subtelomeric sequence of somatic nanochromosomes that encode a single gene, and this is usually within approximately one hundred bases of the transcribed coding sequence (Beh et al. 2019). Since the retained TGLO reading frames are nested within the protein coding sequences of a flanking gene, but also retain their own putative upstream and downstream regulatory sequences, we assessed the expression of retained TGLOs. We collected total RNA from asexually growing cells with the retained TGLO, as well as WT parental lines, and a WT developmental time-course when TGLOs are normally transcribed, and amplified cDNA ends using 5’ RACE (Figure 6A). We found that retained TGLO loci were now transcribed during both the asexual life cycle as well as at their normal developmental pattern (Figure 6B bottom and Figure 6C bottom). The size of the RACE products were similar for the retained lines as well as during normal developmental expression, suggesting that the endogenous TSS was used for gene expression

**Figure 6:**
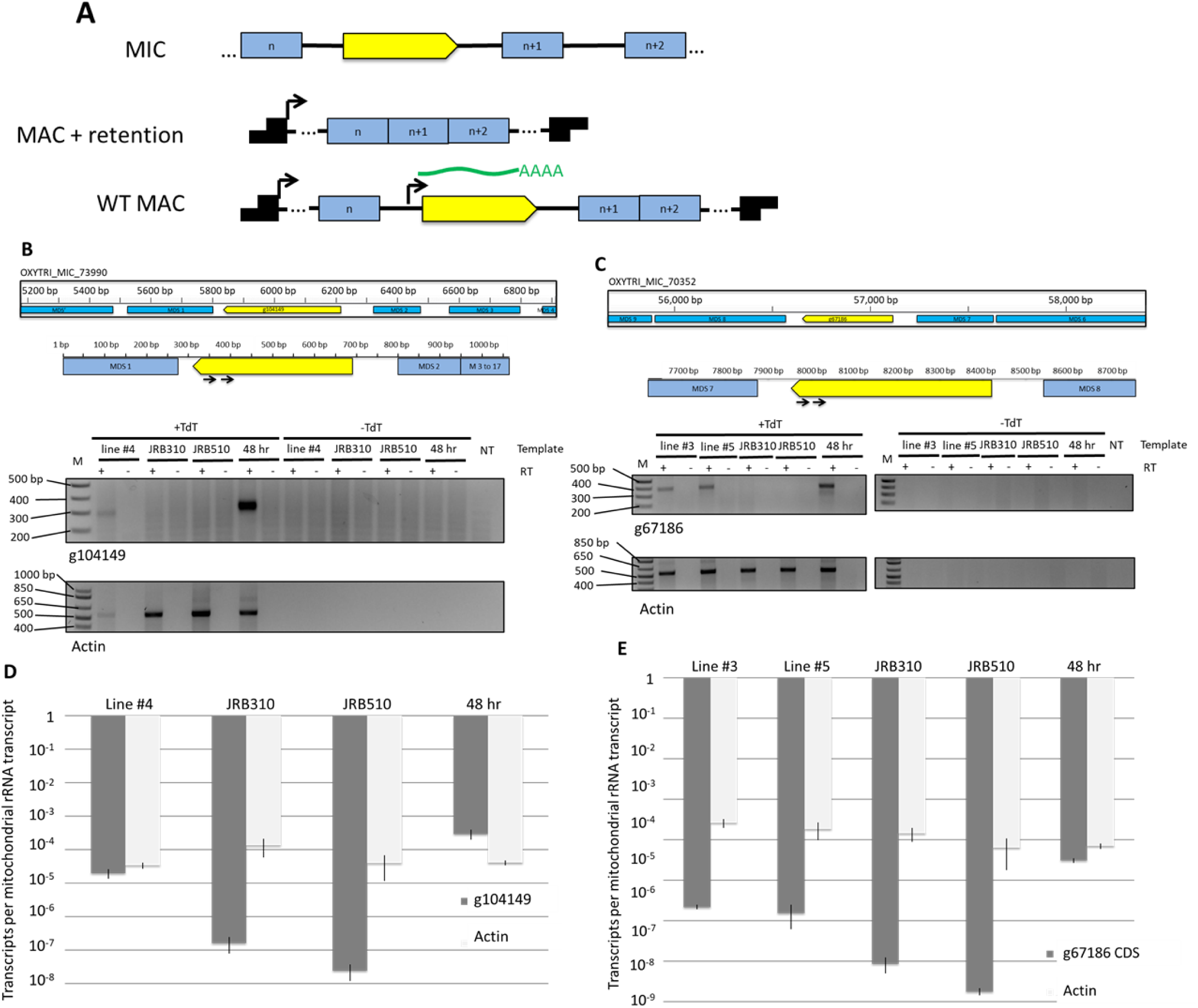
Retained TGLOs are misexpressed during asexual life cycle. **(A)** Possible transcription start sites (black arrows) on a hypothetical rearranged somatic nanochromosome after RNA injection to retain TGLOs (yellow). Green: target transcript deriving from TGLO’s putative upstream regulatory sequence. **(B)** Germline genome locus containing g104149 (yellow) and gene-specific 5’ RACE primers used to amplify transcription start site. **(C)** 5’ RACE products targeting the g104149 or Actin II transcription start site in RNA from F1 retention cells, parental cells, and mid-rearrangement mated cells. TdT: terminal transferase. **(D)** Germline genome locus containing g67186 (yellow) and gene-specific 5’ RACE primers used to amplify transcription start sites. **(E)** 5’ RACE products targeting the g67186 or Actin II transcription start site in RNA from F1 retention cells, parental cells, and mid-rearrangement mated cells. TdT: terminal transferase. **(F)** g104149 or Actin II RNA transcript levels based on qRT-PCR relative to mitochondrial rRNA. Error bars: standard deviation of three biological replicates. **(G)** g67186 or Actin II RNA transcript levels based on qRT-PCR relative to mitochondrial rRNA. Error bars: standard deviation of three biological replicates.

Given the structural differences between the somatic MAC nanochromosome in asexually growing cells and the differentiating MAC during the sexual life cycle, the transcriptional environment of the two nuclei could differ greatly. We used qRT-PCR to compare the transcription levels of retained TGLO loci during the asexual life cycle vs. WT TGLO expression during development, finding that the transcription level of retained TGLOs is approximately an order of magnitude higher during the WT developmental timepoint compared to artificial expression during the asexual life cycle in retained lines (Figures 6D and 6E).

## Discussion

Here, we introduce the definition of TGLO as a transcribed germline-limited DNA sequence with the ability to encode a putative protein. We show that the *O. trifallax* germline MIC genome contains abundant TGLOs that are transcribed to varying degrees in WT cells during development, and are then eliminated from the somatic MAC. This suggests that TGLO gene expression may be regulated by DNA elimination. The conserved domains and predicted functions found in TGLO datasets also support this hypothesis. Moreover, as ciliates have heterochromatic MIC genomes that are not amenable to transcription (Gorovsky and Woodard 1969), and previous observations demonstrated that *Oxytricha*’s germline MIC lacks RNA polymerase II expression (Khurana et al. 2014). Therefore, it is an attractive hypothesis that this lineage may have evolved mechanisms of shutting down gene transcription by programmed DNA elimination after activating gene expression during development.

The earlier report of 810 germline-limited genes in *O. trifallax* assumed that germline-limited coding sequences would be deleted before the cell returned to the asexual life cycle (Chen et al. 2014). Here we present evidence instead that the timing of DNA elimination of TGLOs is heterogeneous during the sexual life cycle. Furthermore, we note the transient addition of *de novo* telomeres in unexpected locations accompanying TGLO transcription, a step that might activate them for transcription. Conceptually similar, in a related ciliate *Euplotes crassus*, DNA processing during the sexual life cycle is responsible for modulating the transcription of one of three telomerase catalytic subunit genes (Karamysheva et al. 2003). Finally, our DNA sequencing results suggest that most TGLOs are indeed eliminated from the somatic MAC by the end of the sexual life cycle. However, we cannot exclude the possibility that a subset of TGLOs persist longer, as further research into later developmental time-points could reveal.

We also observed that at least one germline-encoded ORF, g111288, is actually present at a low somatic copy level in one parental cell line. Unlike TGLOs, g111288 is variably retained as a high copy nanochromosome in some F1 progeny. Presumably, the presence of ncRNAs derived from one parent can program retention of the chromosome in F1 cells, but the incomplete penetrance of somatic g111288 heritability correlates with its low somatic copy number in the JRB510 cell line. Curiously, g111288 does not appear to be transcribed from the somatic MAC in either the parent nor F1 progeny. This is unexpected because the entire coding sequence is present on its own nanochromsome along with its putative upstream and downstream regulatory sequences. However, it is possible that its gene expression requires other *trans-*acting regulatory factors specific to the developmental life cycle.

The case of g111288 is also noteworthy because it appears capable of being either germline-restricted or somatic-encoded. At the level of smaller MDS or IES regions, flexibility between being retained vs. deleted has been observed before but on an evolutionary timescale (Mollenbeck et al. 2006) rather than an intraspecies difference (Vitali et al. 2019). This feature itself could contribute to the birth of new genes, since new coding sequences can sometimes arise from retained noncoding sequences if transcribed and functional (Neme and Tautz 2016; Neme et al. 2017). A previous study in *Tetrahymena* reported that a set of developmentally transcribed somatic minichromosomes are gradually eliminated from the MAC after genome rearrangement (Lin et al. 2016). Moreover, a specific minichromosome in one *Tetrahymena* species might be germline-limited in another species. This *Tetrahymena* example and our functional experiments that reprogram somatic TGLO retention in *O. trifallax* suggest that TGLOs might be a reservoir of sequences with somatic coding potential. We can envision an evolutionary model by which germline-encoded sequences can gain access to the somatic genome where they would be expressed. A deeper intraspecies survey of MAC and MIC genomes, together with developmental RNAseq to survey expression, would be needed to test this hypothesis.

Our ability to program the somatic retention of specific TGLOs via ncRNA injection is a unique feature of the present study. This had the ability to unmask gene expression of targeted TGLOs outside their normal developmental program. *Tetrahymena thermophila* also has non-maintained chromosomes that are lost soon after expression during development and can be fused to adjacent regions to program their retention in the somatic MAC (Feng et al. 2017). Here we have extended this general phenomenon to *Oxytricha* and showed that somatic retention subverts the cell’s endogenous transcription of the gene locus. This supports the hypothesis that TGLO elimination represses their gene expression. In our example the misexpression of a single TGLO locus did not affect cell viability, but the ensemble of loci may need to be silenced during asexual growth.

## Acknowledgements

The authors thank Virginia A. Zakian, Gertrud M. Schüpbach, Samuel Sternberg, and the current and past members of the Landweber lab for their helpful feedback throughout the study. We are grateful to Xiao Chen for bioinformatic help with selecting TGLO candidates. We would also like to thank Wei Wang and Jessica Buckles at the Princeton University Genomics Core Facility for their high-throughput sequencing expertise. This work was funded by NIH R35GM122555 and NSF DMS1764366 to LFL.

